# A Unique LHCE Light-Harvesting protein Family is involved in Photosystem I and II Far-Red Absorption in *Euglena gracilis*

**DOI:** 10.1101/2025.05.07.652572

**Authors:** Héctor Miranda-Astudillo, Rameez Arshad, Félix Vega de Luna, Zhaida Aguilar-Gonzalez, Hadrien Forêt, Tom Feller, Alain Gervasi, Wojciech Nawrocki, Charles Counson, Pierre Morsomme, Hervé Degand, Denis Baurain, Roman Kouřil, Pierre Cardol

**Author notes:** Héctor Miranda-Astudillo, Rameez Arshad, Félix Vega de Luna, Zhaida Aguilar-Gonzalez, Hadrien Forêt, Tom Feller, Alain Gervasi, Wojciech Nawrocki, Charles Counson, Pierre Morsomme, Hervé Degand, Denis Baurain, Roman Kouřil.

## Abstract

Photosynthetic organisms have evolved diverse strategies to adapt to fluctuating light conditions, balancing efficient light capture with photoprotection. In green algae and land plants, this involves specialized light-harvesting complexes (LHCs), non-photochemical quenching, and state transitions driven by dynamic remodeling of antenna proteins associated with Photosystems (PS) I and II. *Euglena gracilis*, a flagellate with a secondary green plastid, represents a distantly related lineage whose light-harvesting regulation remains poorly understood. Although spectral shifts under different light regimes have been observed, their molecular basis was unknown. Here, through integrated phylogenomic, proteomic, structural, and spectroscopic analyses, we identify a novel chlorophyll *a* far-red–absorbing antenna complex in *E. gracilis*, composed of a species-specific Lhce protein family. This antenna forms a pentameric complex under low light and transiently associates with PSII during far-red light exposure. It is structurally and functionally distinct from canonical LHCII□ trimers and absent in Viridiplantae. Additionally, PSI in *E. gracilis* is surrounded by an expanded Lhce/LhcbM belt around a minimal core. These findings reveal a unique mechanism for regulating PS antenna size in *E. gracilis*, distinct from known models in plants and green algae, and highlight an alternative evolutionary strategy for light acclimation in organisms with secondary plastids.

**Highlight:** *Euglena gracilis* features a unique, lineage-specific LhcE antenna system that dynamically associates with PSII and expands PSI light harvesting, revealing an alternative strategy for light acclimation in secondary plastids.

## 1. Introduction

Oxygenic photosynthesis relies on two membrane-integrated protein complexes, Photosystem II (PSII) and Photosystem I (PSI), which work in series to convert light energy into chemical energy. Each photosystem comprises a highly conserved core complex (CC), consisting of a reaction center (RC) and several structural subunits, as well as an internal antenna system. Additional light-harvesting complexes (LHCs) bind to the core complex, capturing and transferring light energy to the reaction center (Cao et al., 2018; Caspy and Nelson, 2018). While PSI and PSII absorb light at different wavelengths, they operate cooperatively during photosynthesis (Emerson, 1957). However, exposure to excessive light can lead to photodamage, particularly in PSII (Aro et al., 1993), though PSI can also be affected under stress conditions such as low temperature (Sonoike, 1995).

PSI contains a limited number of “red” or low-energy chlorophylls, which are critical for regulating energy transfer within its antenna system (Shubin et al., 1992). In land plants, these red chlorophylls are predominantly located in the peripheral light-harvesting complex I (LHCI) (Croce et al., 1998; Croce et al., 2012). In contrast, cyanobacteria, which lack an LHCI antenna, incorporate red chlorophylls directly within their PSI core antenna, with the number and distribution varying among species (Karapetyan et al., 2014). A similar arrangement is observed in *Chlamydomonas reinhardtii*, where the PSI–LHCI supercomplex includes five to six red chlorophylls positioned near the RC, likely at the RC–LHCI interface (Gibasiewicz et al., 2005).

Although the reaction center composition has remained largely conserved throughout the evolution of the primary green lineage (Viridiplantae), the diversity of LHCs has expanded significantly in other photosynthetic eukaryotes (Islam et al., 2020; Koziol et al., 2007; Six et al., 2005). In Viridiplantae, the PSII CC is composed of at least 20 subunits, including the dimeric D1/D2 RC, the inner antenna proteins CP43 and CP47, and over a dozen low-molecular-mass (LMM) transmembrane subunits (PsbE/F/H/I/J/K/L/M/S/Tc/W/X/Z/30/LHCSR). Among these, cytochrome b559 is important for its role in secondary electron transfer pathways, contributing to photoprotection during stress, although it does not participate in primary electron flow (Chu and Chiu, 2016). The oxygen-evolving complex (OEC) includes the extrinsic subunits PsbO, PsbP, and PsbQ. Around the PSII CC are located three minor antenna proteins, Lhcb4 (CP29), Lhcb5 (CP26), and Lhcb6 (CP24, which play critical roles in photoprotection (Graça et al., 2021; Shen et al., 2019). In *Chlamydomonas reinhardtii*, CP29 and CP26 are essential for non-photochemical quenching (NPQ) (Cazzaniga et al., 2020), whereas in land plants, CP24 also participates in NPQ alongside CP29 and CP26 (Miloslavina et al., 2011). Moreover, there is PsbS, which functions as a key light stress sensor in plants and plays a photoprotective role analogous to LHCSR in green algae (Marulanda Valencia and Pandit, 2024; Niyogi and Truong, 2013).

In Viridiplantae, the PSI CC includes the pseudo-symmetric reaction center dimer formed by PsaA and PsaB, ten additional membrane-integrated subunits (PsaF/G/H/I/J/K/L/M/N/O), and a stromal-facing cluster composed of PsaC, PsaD, and PsaE (Naschberger et al., 2022). Among these, PsaG, PsaH, PsaN, and PsaO are unique to green algae and flowering plants. PsaH and PsaO, together with PsaL, contribute to a domain within PSI that binds LHCII during state transitions (Yang et al., 2015). In contrast, red algae contain a PsaO subunit located near the PsaL/A/K interface, while lacking the cyanobacterial-specific PsaX subunit. Although both subunits are unique to their respective lineages, PsaO does not occupy the same position as PsaX and likely serves a distinct structural or regulatory role (Tian et al., 2017; Chen et al., 2022). Several subunits, including PsaA, PsaC, PsaD, PsaE, and PsaF, participate in the interaction with ferredoxin, while PsaF and PsaN mediate plastocyanin binding, completing electron flow through the PSI complex (Amunts et al., 2007; Caspy et al., 2020). *Euglena gracilis* is a photosynthetic flagellate belonging to the Euglenids group, which acquired its complex chloroplast from a primary green alga (Gibbs, 1978; Hallick et al., 1993; Turmel et al., 2009). Unlike primary green plastids, *Euglena* chloroplasts are surrounded by three envelope membranes and lack stacked thylakoid regions (Klein et al., 1972). Our current knowledge of PSII composition in *Euglena gracilis* remains limited, largely due to the lack of comprehensive proteomic and structural data for this organism. The PSII complex in *Euglena* includes core subunits PsbA, PsbB, PsbC, PsbD along with the extrinsic components of the oxygen-evolving complex, PsbO, PsbP, and PsbQ (Suzuki et al., 2004). However, the typical minor antenna proteins CP24 (Lhcb6) and CP26 (Lhcb5) are absent, and only CP29 (Lhcb4) has been detected (Koziol et al., 2007). Additionally, the small subunits PsbX and PsbY, associated with PSII photoinhibition susceptibility (Biswas, 2018), appear to be missing as well (Sobotka et al., 2017). On the PSI side, only five plastid-encoded subunits (PsaA, PsaB, PsaC, PsaJ, and Ycf4) and four nuclear-encoded subunits (PsaD, PsaE, PsaF, and Ycf3) have been identified to date (Sobotka et al., 2017). Notably, most *Euglena* LHCs are synthesized as large polyproteins precursors from mRNAs, and post-translationally cleaved into individual proteins within the chloroplast. Phylogenetic analyses categorized these LHCs into at least eight LhcbM groups (I–VIII) and five Lhca groups (Koziol et al., 2007; Koziol and Durnford, 2008). *Euglena gracilis* possesses pigments of the xanthophyll cycle such as diadinoxanthin and diatoxanthin, but lacks lutein, fucoxanthin, and other pigments typically associated with the violaxanthin cycle. Additionally, its chlorophyll *b* content is relatively low (Cunningham and Schiff, 1986). Light-dependent spectral shifts have been reported (Brown and French, 1961; Winter and Brandt, 1986; Doege et al., 2000), likely reflecting a shared antenna system composed of both LHCI and LHCII proteins that serve both photosystems (Winter and Brandt, 1986; Doege et al., 2000). Notably, *Euglena* appears to lack NPQ as a photoprotective response under excess light (Winter and Brandt, 1986; Doege et al., 2000), and substantial PSII photodamage has been observed under high-light conditions (Doege et al., 2000; Nagao et al., 2021).

In this study, we investigate the structural and functional mechanisms underlying light adaptation in the photosynthetic machinery of *E. gracilis*. Using an integrated phylogenomic, proteomic, and structural approach, we identify a lineage-specific LhcE antenna protein family, reveal its association with both PSI and PSII, and demonstrate its dynamic regulation under far-red and light-limiting conditions. Our findings uncover a novel light acclimation strategy distinct from traditional Viridiplantae mechanisms, involving a far-red–emitting LhcE complex that participates in PSII antenna size dynamic regulation.

## 2. Methods

### 2.1 Algal strain, Growth conditions and Total membrane preparation

*Euglena gracilis* (SAG 1224-5/25) was obtained from the University of Göttingen’s Sammlung von Algenkulturen (Germany). Cells were grown under continuous illumination with a white, fluorescent lamp at photosynthetic photon flux densities (PPFD) of 5, 25, 150, and 500 µmol photons m□² s□¹, corresponding to very low (VLL), low (LL), medium (ML), and high (HL) light conditions, respectively.

The liquid mineral Tris-minimum-phosphate (TMP) medium (pH 7.0) (Harris et al., 2009) was supplemented with a CO_2_ flow (20% in air) and a mix of vitamins (biotin 10^-7^ %, B_12_ vitamin 10^-7^ % and B_1_ vitamin 2 × 10^-5^ % (w/v).

Cells were harvested at the mid-logarithmic phase by centrifugation at 7000 x *g* for 10 min and stored at -70 °C until use. Cell disruption and total membrane preparation by differential centrifugation were performed as described previously (Yadav et al., 2017), with final centrifugation step adjusted to 17000 x *g*. Membrane fractions were stored at -70 °C until further analysis. Protein concentration was determined by the Bradford method (Bio-rad).

### 2.2 Native electrophoresis

Total membrane fractions were solubilized using n-dodecyl-α-D-maltoside (α-DDM) or n-dodecyl-β-D-maltoside (β-DDM) at a detergent-to-protein ratio of 2.0 g/g in solubilization buffer. It contained 50 mM Tris-HCl (pH 8.4), 1.5 mM MgSO_4_, 50 mM aminocaproic acid, 100 mM NaCl, 10% glycerol, 1 mM phenylmethylsulfonyl fluoride (PMSF), and 50 μg/mL tosyl-lysyl chloromethyl ketone (TLCK). The mixture was incubated at 4 °C with gentle agitation for 2 h, and centrifuged at 30,000 x *g* for 30 min.

The resulting supernatants were subjected to high-resolution clear native polyacrylamide gel electrophoresis (*hr*CN-PAGE) (Wittig et al., 2007) using 4%–12% acrylamide gradient gels. To enhance protein resolution, 0.05% sodium deoxycholate and 0.02% α-DM were added to the cathode buffer.

### 2.3 Phylogenetic analyses

The conceptual translations of the complete genomes of 272 organisms broadly sampled across the Tree of Life were downloaded from [<u>figshare Life-OF-Mick</u>]. Those protein sequences were submitted to an orthology inference pipeline relying on NCBI-BLAST v2.2.28+ (Camacho et al., 2009 and OrthoFinder v1.1.2 (Emms and Kelly, 2015)] using an inflation parameter set at 1.2. In parallel, reference sequences for PSI/PSII subunits (and a few other proteins) were compiled from the literature for model organisms (*C. reinhardtii*, *Arabidopsis thaliana*, *Ostreococcus tauri*, *Synechocystis* PCC 6803) (Supplementary Dataset S1). These reference sequences were used to fish out the orthogroups of interest through UBLAST (v7.0.959) (Edgar, 2010) searches. The custom script process-OGs.pl was then used to identify cases of suboptimal delineation among the orthogroups, *i.e.*, when several reference proteins matched a single (possibly quite large) orthogroup and/or when a single reference protein matched several orthogroups. This step led to the combination of pairs or triplets of orthogroups to consolidate families spread over multiple orthogroups. The orthogroups (both single and combined) were then directly aligned with MAFFT v7.273 (Katoh and Standley, 2013). Alignments were manually curated using the editor of the MUST software package v5.60b (Philippe, 1993) and then enriched in orthologous sequences using a prerelease version of Forty-Two v0.211530 (Irisarri et al., 2017; Simion et al., 2017). This allowed us to mine complete genomes and transcriptomes that were not included in the original taxon sampling. We searched four different transcriptomes of *Euglena gracilis*, among which the three public datasets (GDJR01, GEFR01, and HBDM01) analyzed in (Cordoba et al., 2021), to maximize the odds to recover orthologues. Enriched alignments were then filtered to discard too partial sequences and to remove columns containing too many gaps. This was done using ali2phylip.pl (from the Bio-MUST-Core software package (D. Baurain; https://metacpan.org/dist/Bio-MUST-Core)) with the corresponding parameters (min and max) set to 0.50 and 0.01, respectively. Phylogenetic inference was carried out using RAxML v8.1.17 (Stamatakis, 2014) under the model PROTGAMMALG4X and 100 rapid bootstrap replicates. The resulting trees (Supplementary Fig. S1-47) were formatted with format-tree.pl, uploaded to iTOL (Letunic and Bork, 2024) with import-itol.pl and eventually downloaded in their final form with export-itol.pl, all three scripts being also part of Bio-MUST-Core.

For the LHC trees, several additional steps were performed. In particular, the 72 LHC sequences previously reported (Koziol and Durnford, 2008) were supplemented by additional sequences recovered from three transcriptomes (GDJR01, GEFR01, and HBDM01). To this end, the LHC alignment was first stripped out of non-primary green sequences and used in a very sensitive run of Forty-Two (i.e., at a E-value threshold of 1e-05 and with the BRH, trimming, merging, and aligning options all disabled). The 160 candidate transcripts were then submitted to the script xlate-and-splice.pl 1) to assemble the sequences using CAP3 v08/06/13 (Huang and Madan, 1999) (with -p 98 and -o 40), 2) to translate the potential polyproteins in the six possible reading frames and 3) to splice the resulting ORFs based on a PSSM derived from the 44 10-AA linkers described in (Koziol and Durnford 2008). The log L threshold was set to –17 and the minimum ORF/segment length to 24 AA. The 1274 protein fragments obtained were again searched with Forty-Two (setup as above) to recover 165 genuine LHC homologues (out of 100 transcripts). These new sequences were named as follows: <accession>/<frame>/<orf#>/<segment#>@<length>, e.g., GDJR01029061.1+2/F+2/O1/P1@327, then combined with those of Koziol and Durnford (2008 and dereplicated at an identity threshold of 100% with CD-HIT v4.6 (Fu et al., 2012), eventually resulting into 158 unique LHC sequences (Supplementary Dataset S2). The sequences were aligned with MAFFT (linsi), processed with ali2phylip.pl (max = 0.2) and imported into SeaView (Gouy et al., 2010) to build a preliminary tree with PhyML (Guindon et al., 2010) under the LG+F+G4 model; branch support was assessed with aLRT. Based on this tree (Figure S48 and main Figure 1A) and preliminary BLAST and phylogenetic analyses (not shown), final sequence names were attributed, e.g., LHCB4-1_GEFR01021489.1/F-1/O1/P2@205. A second tree (not shown) made of 134 sequences was computed after removing the sequences < 70 AA and those showing long branches in the tree (three sequences were also truncated to remove mispredicted ends). Alignment, filtering, and tree building were carried out as before. The 24 removed sequences are indicated in Supplementary Fig. S48. In parallel, the 134 curated sequences from Euglena were added to the LHC alignment using Two-Scalp v0.243240 (D. Baurain; https://metacpan.org/dist/Bio-MUST-Apps-TwoScalp) (with options linsi and fragments enabled). The enriched alignment was annotated using the reference sequences from Arabidopsis, Chlamydomonas, and Ostreococcus, filtered with ali2phylip.pl (max = 0.4) and submitted to phylogenetic inference using IQ-TREE v1.6.12 (Nguyen et al., 2015) with ModelFinder and ultrafast bootstrap. The resulting tree was rooted, annotated, collapsed, and colored using format-tree.pl, then imported into iToL for further formatting (Figure 1B).

**Figure 1.**
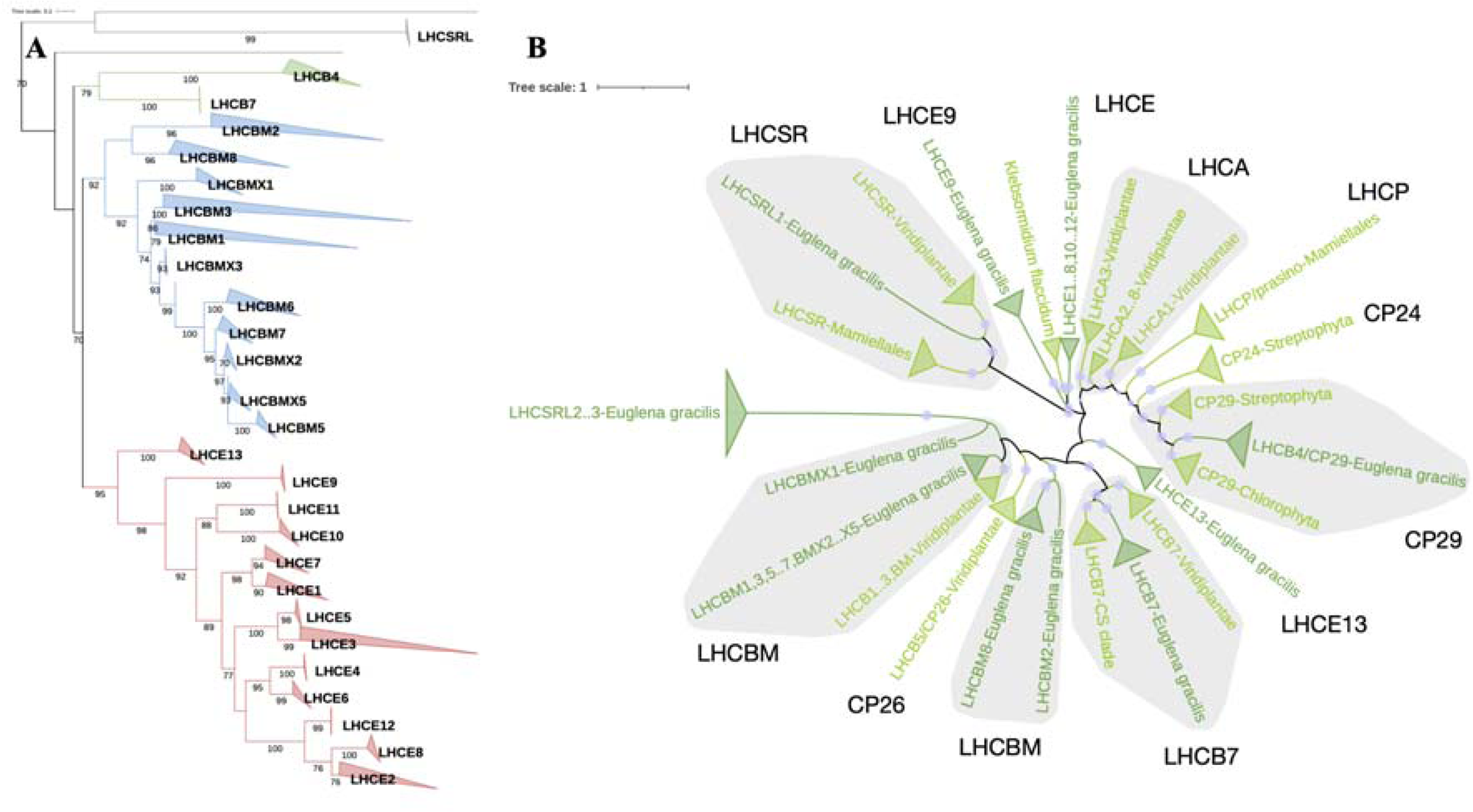
Phylogenetic Analyses of *Euglena gracilis* Light-Harvesting Complex (Lhc) Proteins. (A) **Phylogenetic tree of Lhc proteins in *Euglena gracilis***: The maximum likelihood phylogenetic tree was constructed with PhyML under the LG+F+G4 model using an expanded dataset of *Euglena gracilis* Lhc proteins (158 sequences x 241 sites). aLRT support values ≥70% are indicated. Lhc proteins are grouped into distinct families, with major clades highlighted in different colors: LhcbM proteins (blue), LhcE proteins (red), Lhcb (green) other Lhc-related families (gray). The fully uncollapsed tree showing the position of each individual Lhc sequence (along with its source transcript accession) is available as Supplementary Fig. S48. (B) **Phylogenetic tree of Lhc proteins from Euglena gracilis and Viridiplantae**: The tree was constructed with IQ-TREE under the LG+F+R7 model from an alignment of 405 sequences x 253 sites. It illustrates the evolutionary relationship between *Euglena gracilis* Lhc proteins and those from other Viridiplantae. Collapsed subtrees are all monophyletic and were labelled after the most precise taxon for each group. Ultrafast bootstrap support values ≥70% are shown as light blue bullets. Euglena-specific groups, including LhcbM subfamilies (e.g., LhcbMX1-5) and the newly identified LhcE clade, are highlighted. The presence of distinct Euglena-specific clusters supports the hypothesis of multiple gene duplication events, leading to a unique Lhc composition in *Euglena gracilis*.

### 2.4 Identification of PS subunits

After electrophoretic separation by *hr*CN-PAGE, bands of interest were manually excised from the gel and subjected to liquid chromatography coupled with electrospray-ionization quadrupole time-of-flight mass spectrometry quantitative analysis (LC-ESI-Q-TOF-MS) as previously described (Fox et al., 2020). A single modification was introduced: the mixture underwent reverse-phase chromatography for 35 min. All the obtained sequences were identified using Progenesis software against public protein databases (NRPS/NCBI, UniProt) and a custom in-house database, which is accessible at https://figshare.com/s/57d2ba4ebfbb472ae3de?file=11461148).

LHC proteins-to-PS core ratios were calculated using averaged normalized abundance values from four major PSII core subunits (PsbA, B, C, and D) or PSI core subunits (PsaA, B, D, and F). Two main factors may influence quantification: (i) differential trypsin digestion efficiency across proteins, and (ii) variation in peptide ionization efficiency. Given the sequence and structure similarities between LHCs (Supplementary Fig. S49), we assumed these factors have a negligible impact on LHC quantification. However, variations in abundance of core PSII and PSI subunits (up to a two-fold difference) were observed, prompting us to use averaged values.

### 2.5 Spectrometry

Room temperature absorbance and fluorescence spectra (excitation λ = 470 nm) were obtained using a USB2000+ Ocean Optics spectrometer (Ocean Optics Inc., Dunedin, FL, USA) coupled with a CCD LightBox (Beambio, France). CN gel bands were immediately analyzed after electrophoretic migration.

*In vivo* chlorophyll *a* fluorescence measurements were evaluated with a Joliot Type Spectrophotometer (JTS-10, Biologic, France). Cells were resuspended in fresh medium at a concentration of 5 µg Chl per ml and maintained under shaking dark conditions. Samples were placed into square cuvettes and continuously stirred using a magnetic stirrer. Before fluorescence recording, dark adaptation was conducted for 10 min. Chlorophyll fluorescence was recorded at regular intervals using 10 µs blue detection light pulses. 200 ms saturating pulses of red light (660 nm) were applied to transiently close the PSII reaction centers, allowing measurement of the maximum fluorescence values (Fm). After dark acclimation inside the spectrofluorometer, a weak far-red light (70 µE m^-2^ s^-1^, peaking at 720 nm) was applied for 15 min. Subsequently, both dark-acclimated and far-red-acclimated cells were immediately transferred to a USB2000+ Ocean Optics spectrometer for fluorescence measurements (see above).

The light absorption cross-section of PSII (σ) was determined within the JTS-10 under two different light conditions: red light (25 µE m-2 s-1) and far-red light (300 µE m^-2^ s^-1^). Following dark or far-red light (70 µE m^-2^ s^-1^, peaking at 720 nm) acclimation inside the JTS-10, cells were treated with 10 µM DCMU, an inhibitor of plastoquinone reduction by PSII. After 30 s dark incubation, light was activated, and chlorophyll fluorescence values were recorded. The σ value was calculated as the reciprocal of the time required to reach two-thirds of Fm during fluorescence induction.

### 2.6 Pigment analysis

Bands of interest were manually excised from the CN gel, flash-frozen in liquid nitrogen, and crushed into a fine powder; Pigments were extracted by overnight incubation in 1mL of 100% methanol under vigorous agitation at 4 °C. Following gel extraction, gel debris was removed by centrifugation at 30,000 x *g* for 60 min, and the supernatant was recovered for high-performance liquid chromatography (HPLC) analysis.

Pigments extraction from whole cells was performed as previously described (Gain et al., 2021), and subsequent analysis of HPLC was carried out as previously described (Berne et al., 2018).

### 2.7 Visualization of the isolated Photosystem supercomplexes and LHCE antenna by transmission electron microscopy

Both Photosystem supercomplexes (PSI and PSII) and LHCE bands were eluted from CN gel bands by diffusion in solubilization buffer (see 2.2) supplemented with 0.01% α-DDM. A 4 µl aliquot of each extracted complex solution was applied on freshly glow-discharged, carbon-coated copper grids. Excess sample was blotted with filter paper and the grids were negatively stained with 2% uranyl acetate to enhance contrast.

Imaging was performed using a Tecnai T20 transmission electron microscope equipped with a Gatan 4000 SP 4K slow-scan CCD camera. Automated data acquisition was conducted to capture images of 2048×2048 pixels, with a magnification of 133,000x magnification and a pixel size of 0.225 nm. A total of 8,034, 7,615 and 3350 micrographs were recorded for PSII, PSI and LHCE samples, respectively.

From the selected micrographs, 194,090, 85,435 and 3,840 single particles of PSII, PSI and LHCE complexes, respectively, were independently picked and subjected to reference-free 2D alignment and classification using the image processing framework SCIPION (de la Rosa-Trevín et al., 2016). Structural models were generated by fitting 2D projection maps with high-resolution structural models obtained from Protein Data Bank: PSII from *Chlamydomonas reinhardtii* (PDB: 6KAD) (Sheng et al., 2019) and PSI from *Dunaliella salina* (PDB 6RHZ) (Perez-Boerema et al., 2020).

## 3. Results

### 3.1 Expanding the Phylogenetic Landscape of Euglena gracilis Lhc Proteins: Evolutionary Insights and Novel Subfamilies

To investigate the phylogenetic distribution and diversity of Lhc proteins in *Euglena gracilis*, we expanded the existing dataset (Koziol and Durnford, 2008; Koziol et al., 2007) by incorporating sequences from four recently released *E. gracilis* transcriptomes (Cordoba et al., 2021). This resulted in a comprehensive set of 158 predicted Lhc proteins (Supplementary Fig. S48 and supplementary Dataset S2). The expansion in sequence number is likely attributable to the presence of polyprotein precursors and extensive gene duplication (Koziol and Durnford, 2008), as well as the proposed triploid nature of the *E. gracilis* nuclear genome (Fields et al., 2024).

Using an updated and broader reference set of Viridiplantae genomes compared to previous study (Koziol et al., 2007), we constructed a phylogenetic tree in which *E. gracilis* Lhc proteins were classified into over 20 distinct Lhc families (Figure 1A; Supplementary Fig. S48). While the LhcbM family had previously been divided into eight major subfamilies (Koziol and Durnford, 2008), our expanded analysis reveals that most *Euglena* LhcbM sequences form a sister group to the Viridiplantae LhcbM family. Importantly, we identified five additional Euglena-specific groups, designated LhcbMX1 to LhcbMX5, not reported in earlier studies (Koziol et al., 2007; Koziol and Durnford, 2008).

Three subgroups displayed especially divergent evolutionary positions with no clear orthologs in other species: LhcbM2 and LhcbM8 clustered on a basal branch of the Lhcb subtree, while LhcbM4 (renamed here LhcE13) family also formed an independent cluster. Additionally, none of the so-called *Euglena* Lhca proteins grouped with either Lhcb or Lhca families from green algae or land plants (Figure 1B), indicating that these sequences represent a separate evolutionary lineage. This pattern suggests that the *E. gracilis* Lhc protein family has undergone extensive lineage-specific diversification, likely via multiple gene duplication events from a limited ancestral Lhc genes, a trajectory reminiscent of the LhcP expansion in Prasinophyceae (Six et al., 2005) and the Lhcf family in diatoms (Islam et al., 2020). Based on these findings, we propose a new protein family designation, LhcE, where “E” stands for *Euglena* or *Euglenozoa*. A full nomenclature reference is provided in Supplementary Table S1).

Unlike LhcbM proteins, all members of the proposed LhcE family, including the renamed LhcE13 (formerly LhcbM4), lack the canonical trimerization motif WYGP(D)R (Koziol et al., 2007; Koziol and Durnford, 2008), further supporting their functional and structural divergence. Furthermore, our analysis did not identify any sequences corresponding to the PSII minor antenna proteins CP26 (Lhcb5) and CP24 (Lhcb6), confirming earlier observations (Koziol and Durnford, 2008). Only orthologs of CP29 (Lhcb4) and Lhcb7 were detected, underscoring the distinctive Lhc composition of *E. gracilis*.

### 3.2 Identification and Characterization of Photosynthetic Complexes Associated with Euglena gracilis Lhc Proteins

To identify the photosynthetic complexes that associate with *Euglena gracilis* Lhc proteins, total membranes from photoautotrophically grown cells were solubilized using either *n*-dodecyl-α-D-maltoside (α-DDM) or *n*-dodecyl-β-D-maltoside (β-DDM). The resulting pigment–protein complexes were separated by high-resolution clear native electrophoresis (hrCN-PAGE). This analysis revealed eight and six major green bands for α-DDM and β-DDM extracts, respectively, with apparent molecular weights ranging from 110 to 1,500 kDa (Figure 2A; Supplementary Fig. S50). Since α-DDM allowed the solubilization of complexes with higher molecular weight than β-DDM, we focused on α-DDM solubilized material for subsequent characterization.

**Figure 2.**
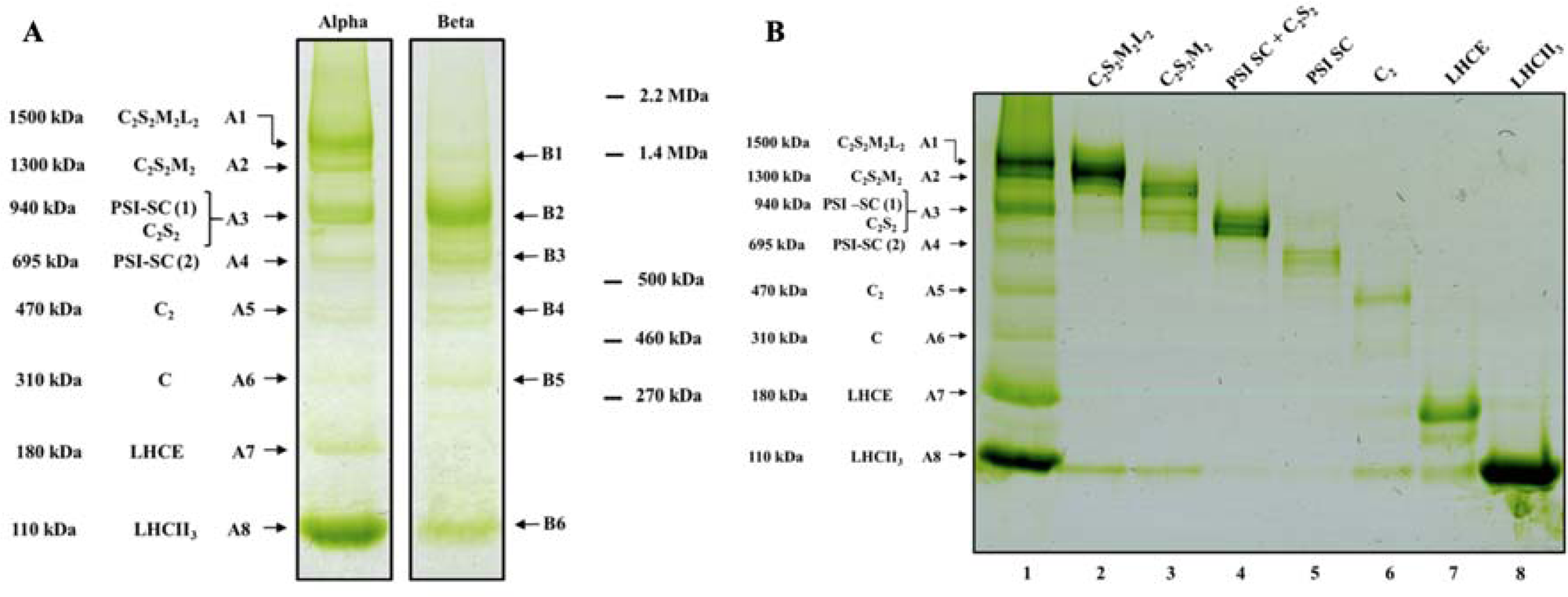
A. Comparison of pigment-protein complexes extracted using two mild detergents and analyzed by hrCN-PAGE. Total membrane extracts *from Euglena gracilis* were solubilized with n-dodecyl-α-D-maltoside (α-DDM) and n-dodecyl-β-D-maltoside (β-DDM), followed by high-resolution clear native polyacrylamide gel electrophoresis (hrCN-PAGE). α-DDM solubilization yielded eight major chlorophyll-containing bands (A1–A8), ranging from 110 to 1500 kDa. β-DDM solubilization resulted in six major bands (B1–B6), spanning 310 kDa to 2.2 MDa. Identified photosynthetic complexes include PSII-LHC supercomplexes (C□S□M□L□, C□S□M□, C□S□), PSI supercomplexes (PSI-SC), PSII core (C□), the LHCE antenna complex, and free LHCII□ trimers. These results highlight differences in extraction efficiency between α-DDM and β-DDM, with α-DDM favoring the solubilization of larger supercomplexes, and LHCE antenna complex stability. **B. *hr*CN-PAGE of purified photosynthetic pigment-proteins complexes**. Each band from α-DDM extracted membrane samples was manually excised, extracted under liquid nitrogen, and analyzed by liquid chromatography-electrospray-ionization quadrupole time-of-flight mass spectrometry (LC-ESI-Q-TOF-MS) (see Methods 2.3 and 2.5 for details). A sample of each purified complex was reloaded onto lanes 2–8 for validation. Lane 1, total membrane extract before purification; lanes 2–8, purified complexes corresponding to distinct photosynthetic assemblies. The purified complexes display distinct migration patterns, confirming their identity and integrity after extraction and analysis.

The hrCN-PAGE bands (A1–A8) from α-DDM-solubilized samples were excised and subjected to two complementary analyses: (i) reloading onto a second hrCN-PAGE gel to assess their stability and possible subcomplex dissociation (Figure 2B), and (ii) protein identification by liquid chromatography–electrospray ionization–quadrupole time-of-flight mass spectrometry (LC-ESI-Q-TOF-MS) (Supplementary Table S2). As the second hrCN-PAGE separation shows, the two largest complexes (bands A1 and A2; ∼1.5 and ∼1.3 MDa, respectively) predominantly contained PSII core subunits along with LhcbM proteins. The next two complexes (bands A3 and A4; ∼940 and ∼695 kDa) were enriched in PSI core subunits, as well as LhcE proteins and some LhcbMs. The presence of PSII subunits in band A3 suggests possible co-migration of PSI and PSII particles of similar size. Complexes of intermediate molecular weight (bands A5 and A6; ∼470 and ∼310 kDa) consisted mainly of PSII core components, with a limited amount of bound LHCs. The smallest complexes (bands A7 and A8; ∼180 and ∼110 kDa) were composed exclusively of LhcE and LhcbM proteins, respectively. Interestingly, the PSII-enriched high-molecular-weight complexes (A1 and A2) released the smaller LhcbM-only complex (A8) during purification (Figure 2B, lanes 2 and 3). This observation supports the hypothesis that band A8 components are originally integrated into the larger PSII–LhcbM supercomplexes and may dissociate during detergent extraction or purification.

### 3.3 CP26-Less PSII Core Associates with Up to Three LHCII Trimers

Through integrated phylogenomic and proteomic analyses of separated gel bands A1-A8, we identified orthologs of a broad set of PSII subunits in *Euglena gracilis*, including PsbA–F, PsbH–R, PsbT, PsbW–Z, and Psb27, Psb28, Psb29, Psb32, and Psb33 (Supplementary Table S3, and supplementary Dataset S3). This significantly expands the list of known PSII subunits in *Euglena* compared to an earlier report (Sobotka et al., 2017). Notably, we did not detect sequences corresponding to PsbS (supplementary Fig. S19), a key light-stress sensor involved in photoprotective mechanisms in plants, or to PsbU and PsbV (supplementary Fig. S21-22), which are typically found in cyanobacteria and red algae. Overall, the PSII core composition in *E. gracilis* closely resembles that of green lineage model organisms such as *Chlamydomonas reinhardtii* (Shen et al., 2019) and *Arabidopsis thaliana* (Graça et al., 2021).

In the high-molecular-weight PSII–LHC supercomplexes (bands A1 and A2), the associated LHC proteins include CP29 and LhcbM isoforms from the LhcbM1, LhcbM5, LhcbM6, and LhcbM7 protein families (Figure 3A). These same LhcbM isoforms were also detected in the ∼110 kDa complex (band A8), which likely corresponds to free LHCII trimers (LHCII□) (Figure 3A). In contrast, LhcbM proteins were absent from the intermediate-mass PSII complexes at 470 and 310 kDa (bands A5 and A6), which we tentatively identify as related to dimeric (C□) and monomeric (C□) PSII cores, respectively (Figure 3A, and Supplementary Table S2).

**Figure 3.**
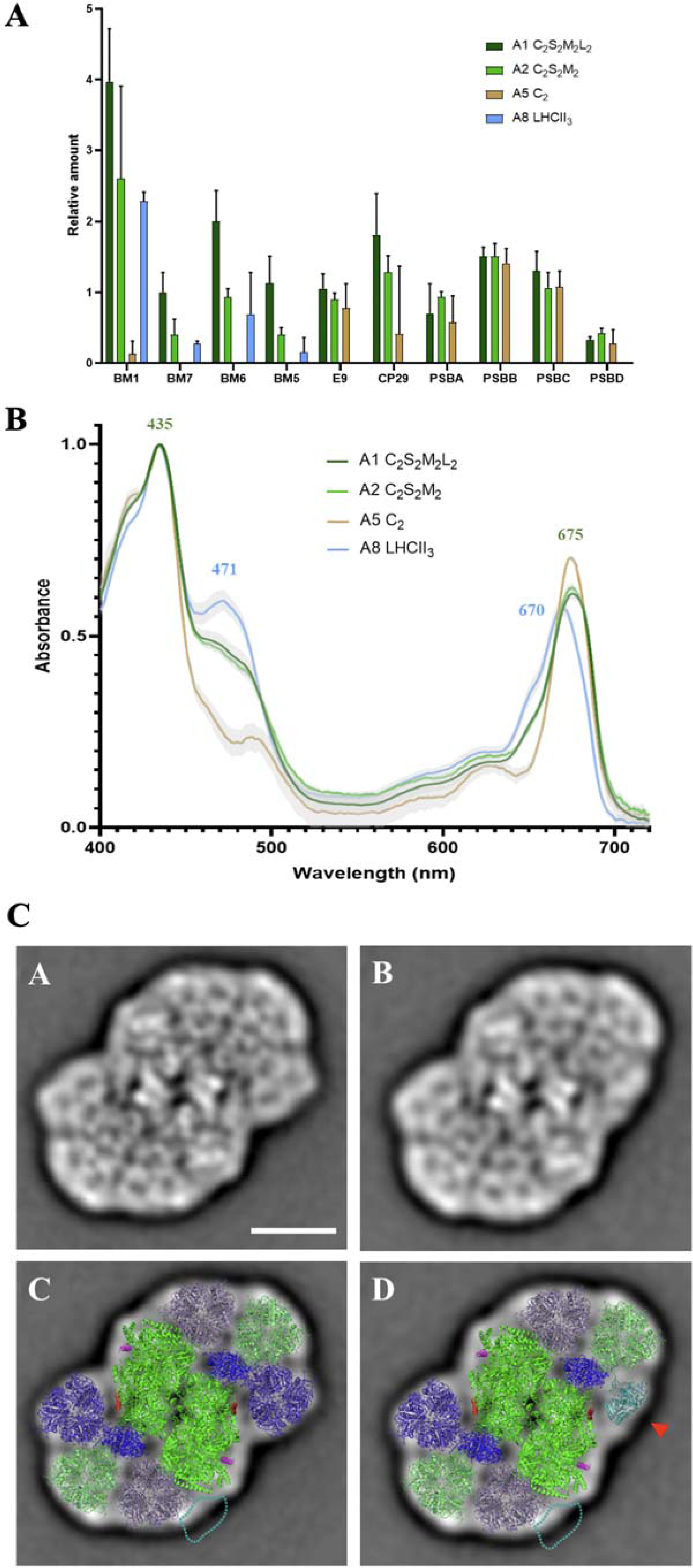
*E. gracilis* photosystem II. **A Quantification of LHC proteins associated with PSII supercomplexes, PSII core, and free LHCII□.** Relative amounts of different LhcM subtypes (LhcbM-1, 5, 6, 7, and CP29) and LhcE9 (E9) were quantified using a proteomic approach across various PSII complexes. The four PSII complexes analyzed include C□S□M□L□ (dark green), C□S□M□ (light green), C□ (brown), and free LHCII□ (blue). Spectral count assigned to each antenna was normalized to the average spectral count assigned to the core complex polypeptides (calculated as (PsbA+PsbB+PsbC+PsbD)/4) in the same band. Relative number of LhcbBM proteins in free LHCII trimer is calculated assuming a total of 3 LHCS. Error bars represent standard deviations from 3 independent biological replicates. **B**. **Absorption spectra of PSII complexes LHCII trimers.** Absorption spectra for bands 1, 2, 5, 7 and 8, normalized to their maximum absorbance. Mean values are represented as solid lines, with standard deviations shown as shaded areas (n=3 based on independent biological replicates). The different PSII and antenna complexes are color-coded as follows: C□S□M□L□ (complete PSII-LHC supercomplex) in blue; C□S□M□ (PSII-LHC intermediate complex) in yellow green; C□ (PSII core complex) in green; LHCII□ free trimer in red. **C**. Projection maps and structural models of PSII supercomplexes from *Euglena gracilis* revealed by single particle electron microscopy. (A-B) Projection maps of two forms of the largest PSII supercomplex (C_2_S_2_M_2_L_2_). (C-D) Structural models of different forms of the PSII supercomplexes obtained by fitting the high-resolution structure of the PSII supercomplex from green alga *Chlamydomonas reinhardtii* (PDB: 6AKD (Shen et al., 2019). (C) The larger form consists of a dimeric PSII core complex (green), minor antenna proteins CP29 (blue), and three pairs of LHCII trimers. (D) The smaller forms lack one L-trimer (red arrow). The reduced density reflects a PSII heterogeneity (see text for details). The scale bar is 10 nm.

To estimate the stoichiometry of antenna proteins in each complex, we calculated the LhcbM-to-core ratios based on the relative band intensities of four major PSII core subunits (PsbA/B/C/D) as shown in Figure 3A. The 1.5 MDa supercomplex displayed an LhcbM-to-core ratio of 8.5, while the 1.3 MDa complex showed a ratio of 4.4. Importantly, the relative proportions of the four LhcbM types were nearly identical between the PSII supercomplexes and the 110 kDa LhcbM-only complex (Figure 3A; Supplementary Fig. S51), reinforcing the idea that the latter represents LHCII□ dissociated from the PSII-SC antenna. In addition, CP29 (Lhcb4) and the *Euglena*-specific LhcE9 protein were consistently associated with the PSII supercomplexes at an apparent 1:1 stoichiometry, suggesting a stable integration into the antenna structure (Figure 3A).

#### PSII-SC Structure and LHCII□ Association

Single-particle EM analysis of the 1.5 MDa *E. gracilis* PSII-SCs (band A1) yielded 2D projection maps revealing a dimeric PSII core, with each monomer associated with up to ten LHC proteins, nine of which are arranged into three LHCII_3_ (Figure 3C). This organization is structurally similar to the C□S□M□L□-type PSII-SC previously characterized in *Chlamydomonas reinhardtii* under similar solubilization conditions (Burton-Smith et al., 2019; Shen et al., 2019). Based on this structural similarity and the molecular mass, the 1.3 MDa complex likely corresponds to C_2_S_2_M_2_.

The structural identification of a C□S□M□L□-type PSII-SC in *E. gracilis* is consistent with spectroscopic and pigment data, which also point to a high LHCII content. Its chlorophyll *a/b* ratio (Table 1) and relative absorbance at 480 nm (Soret band) are higher compared to the PSII core complex, but remain lower than the values observed for the free LHCII trimers (Figure 3B). Variations in the number of LHCII□ trimers associated with C_2_ were also evident from EM analysis of smaller PSII-CSs (band A3) (Supplementary Fig. S52). These data suggest a dynamic PSII antenna system in *E. gracilis*, capable of forming supercomplexes with variable LHCII stoichiometry. Such structural variation is also characteristic of land plants (Kouřil et al. 2018), where PSII supercomplex heterogeneity contributes to light adaptation and energy distribution.

**Table 1.** Pigment composition of pigment-protein complexes of *E gracilis*.

| Pigment | C<sub>2</sub>S<sub>2</sub>M<sub>2</sub>L<sub>2</sub> | C<sub>2</sub>S<sub>2</sub>M<sub>2</sub> | PSI-SC | PSI-SC | C<sub>2</sub> | LHCE | LHCII<sub>3</sub> |
| --- | --- | --- | --- | --- | --- | --- | --- |
| Neoxanthin | 1.4 ± 0.02 | 2.0 ± 0.06 | 0.7 ± 0.04 | 0.8 ± 0.05 | 3.0 ± 0.2 | 0.5 ± 0.01 | 9.1 ± 0.1 |
| Diadinoxanthin | 10.9 ± 0.3 | 8.3 ± 0.2 | 13.0 ± 0.8 | 10.9 ± 0.8 | 15.6 ± 1.2 | 18.1 ± 0.4 | 20.2 ± 0.3 |
| Diatoxanthin | 0.8 ± 0.03 | 0.9 ± 0.02 | 1.1 ± 0.1 | 0.6 ± 0.03 | 0.8 ± 0.2 | 0.8 ± 0.1 | 1.7 ± 0.03 |
| Chlorophyll *b* | 20.8 ± 0.6 | 18.9 ± 0.5 | 7.4 ± 0.4 | 5.4 ± 0.3 | 13.1 ± 1.6 | 2.6 ± 0.13 | 27.7 ± 0.3 |
| Chlorophyll *a* | 65.3 ± 1.6 | 69.2 ± 1.8 | 77.2 ± 5.2 | 81.7 ± 5.7 | 66.6 ± 5.1 | 77.2 ± 1.5 | 40.6 ± 0.5 |
| Beta-carotene | 0.8 ± 0.14 | 0.67 ± 0.02 | 0.65 ± 0.06 | 0.61 ± 0.03 | 0.84 ± 0.02 | 0.75 ± 0.27 | 0.64 ± 0.01 |
| *a/b* ratio | 3.1 | 3.7 | 10.4 | 15.2 | 5.1 | 29.8 | 1.5 |

Data represent mean ± standard deviation (s.d.) of the relative mass of each pigment (n = 3, based on independent biological replicates).

#### Absence of CP26 and Potential Functional Implications

EM analysis confirmed that CP29 is consistently associated with the *E. gracilis* PSII core complex, while CP26 (Lhcb5), a conserved minor antenna typically positioned adjacent to the S-trimer in *Chlamydomonas* PSII-SC (Burton-Smith et al., 2019; Shen et al., 2019), is absent and not replaced by any other antenna subunit (Figure 3C). This structural finding agrees with our phylogenomic analysis (Section 3.1), which revealed no CP26 ortholog among the 158 identified *E. gracilis* LHC proteins.

EM projections also suggest PSII heterogeneity. In several views, one of the L-trimers appears incomplete (Figure 3C, panels B and D), raising the possibility of structural variation within PSII-SCs. One hypothesis is that the missing LHCII subunits are replaced by a monomeric antenna protein, potentially LhcE9, which was consistently detected in PSII-SCs by proteomic analysis (Figure 3A). Such substitution may represent a novel mechanism for modulating PSII antenna size and composition in response to fluctuating light conditions. Alternatively, it could be a partial disintegration of the L-trimer, with only two monomeric LhcbM subunits remaining attached to the core.

### 3.4 Minimal PSI Core Is Surrounded by an LhcE/LhcbM Belt

Our phylogenomic analysis confirmed earlier findings (Sobotka et al., 2017) regarding the presence of PsaF and PsaJ and the absence of PsaI, PsaK, and PsaL in *Euglena gracilis* (Supplementary Table S3; Supplementary dataset S3). By extending our analysis to additional PSI subunits typically found in the green lineage (Scheller et al., 2001), we identified PsaM as a chloroplast-encoded component (Supplementary Fig. 45), while no orthologs of PsaG, PsaH, PsaN, or PsaO were detected (Supplementary Fig. S40-41 and S46-47).

In land plants, PsaG and PsaK play important roles in anchoring LHCI to the PSI core (Huang et al., 2021; Ozawa et al., 2018). Their absence in *E. gracilis* raises questions about how the PSI–LHC supercomplex is structurally organized in this organism. Mass spectrometry–based quantification revealed an average of 10 Lhc proteins per PSI core in the 940 kDa complex (band A3), comprising primarily LhcE subunits (LhcE5–8, LhcE10–11, and LhcE13), along with 2–3 LhcbM proteins (LhcbM2 and LhcbM8) (Figure 4A). The fact that these LHCBM types were not detected in PSII supercomplexes or in the free LHCII trimer by mass spectrometry (Figure 3A, Supplementary Table S2) suggests that they are either not functionally associated with PSII or represent a distinct pool of LHC proteins, possibly dedicated to other assemblies such as PSI or peripheral antenna complexes.

**Figure 4.**
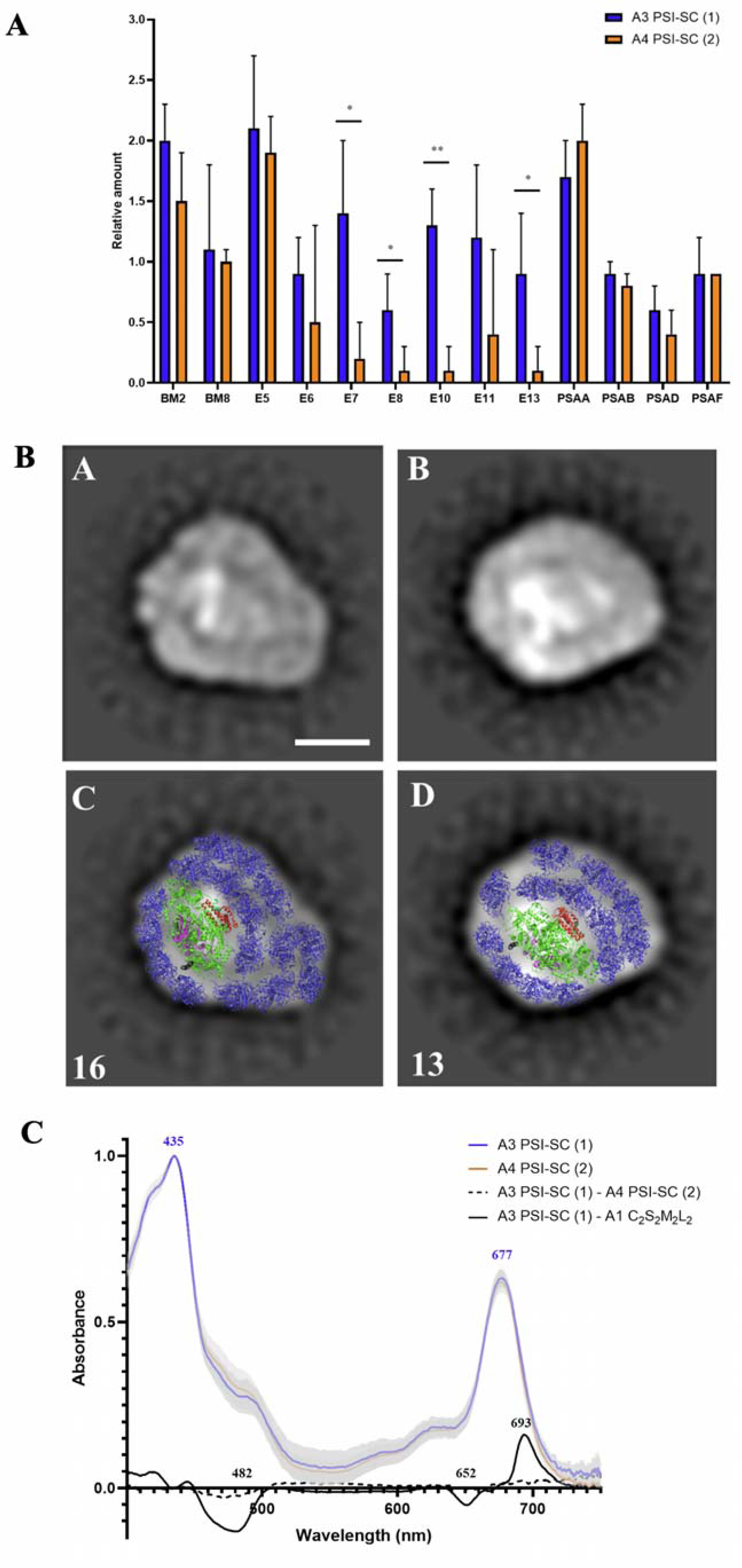
*E. gracilis* photosystem I supercomplex. **A. Antenna composition of PSI–LHC supercomplexes determined by MS/MS analysis.** Lhc proteins from two different branches are present (see Figure 1 and Supplementary Fig. S48 for details): LhcE5, E6, E7, E8, E10, E11, E13, bM2, and bM8. Spectral count assigned to each antenna was normalized to the average spectral count assigned to the core complex polypeptides (calculated as (PsaA+PsaB+PsaD+PsaF)/4) in the same band. Error bars represent standard deviations from 3 independent biological replicates (A3) and 4 independent biological replicates (A4). Statistical analysis was performed using an independent unpaired t-test. Asterisks indicate statistically significant differences (*, *p* < 0.05; **, *p* < 0.005). **B. Structural models of PSI-LHCI supercomplexes revealed by single-particle electron microscopy**. Structural models of different PSI-LHCI supercomplex forms obtained through single-particle electron microscopy. Top rows: raw electron density projections. Bottom rows: corresponding fitted models with assigned structural components. Densities were assigned by fitting the high-resolution PSI structure from *Dunaliella salina* (PDB: 6RHZ, Perez-Boerema et al., 2020). Additional antenna proteins were modeled using copies of the Lhca1 subunit from *Dunaliella salina*. PSI core (green), PsaD subunit (violet); LHC (blue) belt at the PsaF side (red); Variable “flipper” region at the PsaM side (black) (multicolored LHC subunits). The structural variability observed highlights differences in antenna organization, providing insights into the dynamic nature of PSI light-harvesting adaptations. The scale bar represents 10 nm. **C. Absorption spectra of PSI-LHC complexes**. Absorption spectra for bands 3 and 4, corresponding to PSI-LHC complexes, normalized to their maximum absorbance. Mean values are shown as solid lines, with standard deviations represented as shaded areas (n=3, based on independent biological replicates). PSI-LHC (band A3 in Figure 1A-B) in blue, and PSI-LHC (band A4 in Figure 1-AB) in orange. These spectra highlight the similarity in light absorption profiles between the two PSI-LHC complexes, with distinct peaks in the visible (∼440 nm) and far-red (∼680 nm) regions, characteristic of PSI-bound antenna systems.

Single-particle EM analysis of the 940 kDa PSI-SC (band A3) revealed its raw architecture in which a minimal PSI core, composed of PsaA, PsaB, PsaC, PsaD, PsaE, PsaF, PsaJ, and PsaM can be fitted and is surrounded by rows of antenna proteins, comprising a total of 11 to 16 Lhc subunits (Figure 4B, Supplementary Fig. S53). An unusual antenna arrangement is proposed: on the PsaM side (shown in black), a compact and stable Lhc belt could accommodate four tightly associated Lhc proteins, resembling the conserved LHCI configuration found in the PSI–LHCI SCs of land plants (Wang et al., 2021) or the first LHCI row in *C. reinhardtii* (Huang et al., 2021).

In contrast, a broad band of Lhc proteins lines the side of the PsaF core (shown in red), forming an elongated structure that wraps toward the PsaA/B boundary and creates a variable “flipper” region in the outer domain of the antenna. This region appears to include Lhc subunits forming a second, and in some cases even a partial third, antenna layer. A similar organization of the PSI antenna system has been described in prasinophyte green algae, where four lineage-specific LHCPs are located on the PsaM side opposite the conserved PsaF-LHCI belt, although these lack the far-red absorption features observed in *Euglena* (Swingley et al., 2010). In moss (*Physcomitrella patens*), PSI SCs include nine LHCI proteins and one LHCII□ unit; however, the antenna extension occurs on the PsaK side, in contrast to *Euglena*, where expansion is observed on the PsaM side (Iwai et al., 2018, Pinnola et al. 2018). Since we also detected approximately three LhcbM proteins per PSI core (Figure 4A), we also attempted to model the presence of the LHCII□ trimeric unit within the PSI supercomplex. However, due to the limited resolution of the projection maps, no clear density corresponding to LHCII□ was observed. This suggests that any association of LHCII□ with PSI may be transient or structurally labile, potentially lost during sample preparation for EM analysis.

Biochemical comparison of the larger and smaller PSI supercomplexes (bands A3 and A4, respectively) indicates that the lower molecular weight form preferentially lacks LhcE7, LhcE8, LhcE10, and LhcE13 (Figure 4A). This suggests that these subunits are part of a more peripheral or weakly bound antenna region, likely within the flexible flipper zone, as inferred from projection maps of the smaller PSI supercomplex (Figure 4B, panels B and D, and Supplementary Fig. S53). Despite these differences in antenna size, both PSI–LHC assemblies exhibit similar spectral features, with a main absorption peak at 677 nm and a prominent far-red shoulder at 693 nm (Figure 4C), distinguishing them from PSII–LHC complexes.

### 3.5 An LhcE Antenna Complex Accounts for Far-Red Absorption in Euglena gracilis cells under Low-Light Conditions

The 180 kDa LhcE antenna complex (band A7) primarily contains diadinoxanthin and chlorophyll *a* (Table 1). As a result, its absorption spectrum lacks the characteristic chlorophyll *b* peaks at 480 and 653 nm. While LhcbM trimers (band A8) show a maximum absorbance at 672 nm, the LhcE complex displays red-shifted absorption peaks at 675 and 692 nm (Figure 5A). At room temperature, its fluorescence emission is also red shifted, with a maximum at 698 nm, compared to 684 nm for LHCII□ (Figure 5B). A red-shifted antenna complex in *E. gracilis* was previously reported (Doege et al., 2000), but no further characterization had been performed until now.

**Figure 5.**
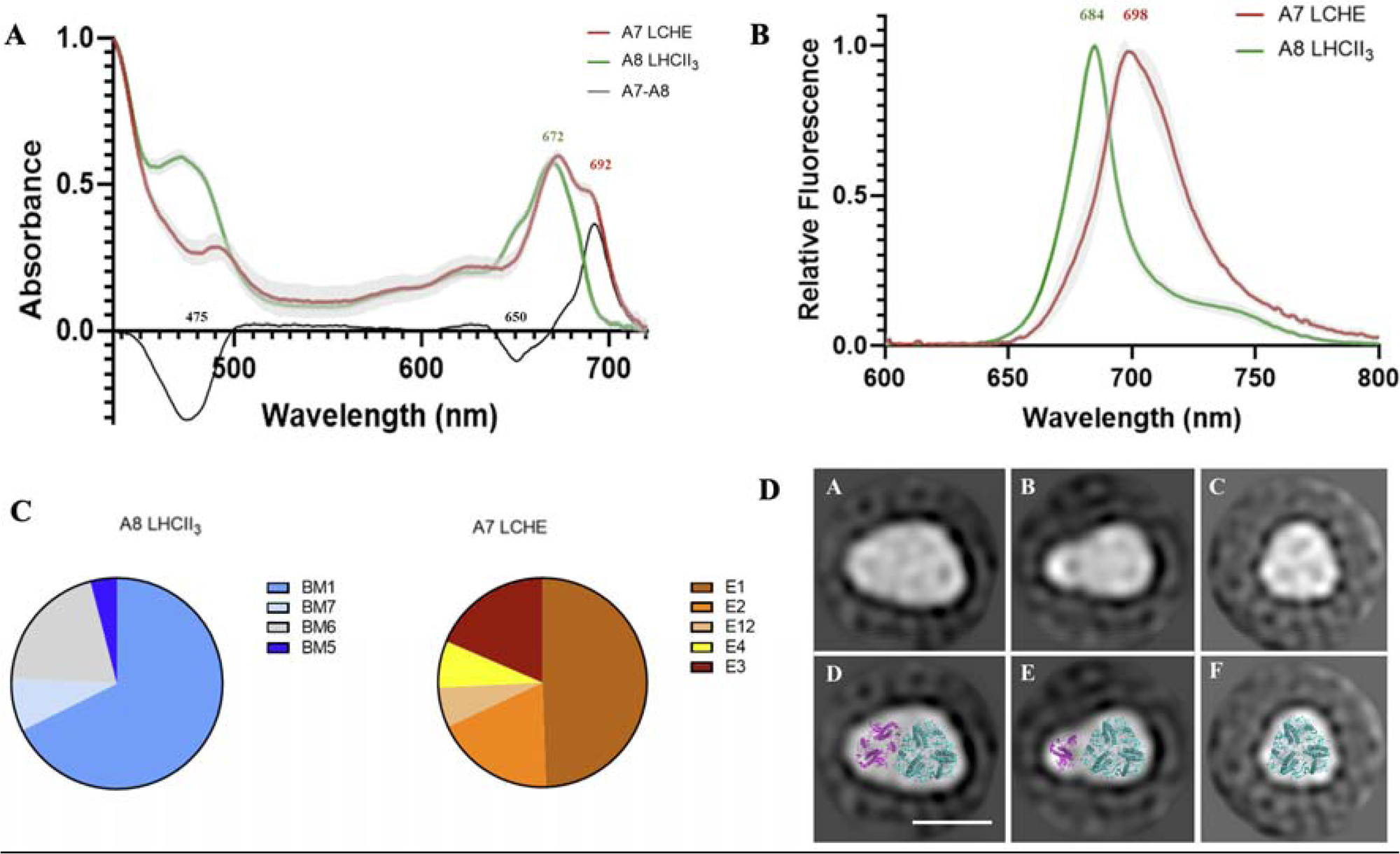
LCHE antenna complex. **A**. Absorption spectra of isolated LHCE antenna complex and LHCII trimers. **B.** Room temperature fluorescence spectra of isolated LHCE antenna complex and LHCII trimers. **C.** Relative amounts of LHC proteins in the 110 kDa (LHCII_3_) and 180 kDa (LHCE_5_) antenna complexes. (n=3, based on independent biological replicates) **D.** Averaged 2D projections of the LHCE antenna, three major classes were collected with a class sum of 4028, 2907, and 3840 particles, for A, B, and C, respectively. The larger LHCE complex comprises up to 5 LHCE monomers (A and D): 1 trimer (cyan) + 2 monomers (violet). Smaller classes may correspond to LhcE antenna complexes with detached monomers (Panels B, C, E and F). The scale bar is 10 nm.

Based on its molecular mass, the 180 kDa complex likely comprises five to six LHC proteins, among which LhcE1, LhcE2, LhcE3, LhcE4, and LhcE12 were identified in varying proportions by mass spectrometry (Figure 5C). Notably, these LhcE proteins were not detected in association with PSI– LHC or PSII–LHC supercomplexes (Figures 3A and 4A), suggesting that the LhcE antenna complex is peripheral and readily dissociates during α-DDM solubilization. When β-DDM was used instead of α-DDM, the LhcE complex was destabilized—like the largest PSII–LHC supercomplex (Figure 2A), indicating weaker binding affinity or structural sensitivity to detergent conditions.

Single-particle EM analysis of the 180 kDa complex identified three main projection classes (Figure 5D). In the largest projection, five LhcE proteins could be fitted. The two smaller projections likely correspond to subcomplexes derived from partial disassembly of the pentameric antenna, although the smallest projection may also reflect contamination from LhcbM trimers. Due to the structural similarity of these projections to LHCII□ in PSII supercomplexes (Figure 3E), but despite lack of trimerization motif for LHCE proteins (Koziol et al., 2007; Koziol and Durnford, 2008), we propose that the 180 kDa complex contains one LhcE trimer along with additional monomeric subunits (Figure 5D). This interpretation is consistent with the hrCN-PAGE separation profile of the LhcE fraction, in which lower molecular weight subcomplexes appear as distinct bands (Figure 2B, lane 7).

Interestingly, the difference absorption spectrum between the LhcE complex and LhcbM trimers (Figure 5A) closely mirrors the difference between PSI–LHC and PSII–LHC SCs (Figure 4C). In both cases, the spectra show negative peaks at 470–480 nm and 650–652 nm, corresponding to the absence of chlorophyll *b*, and a maximum at 692–693 nm, indicative of enhanced far-red absorption by red-shifted chlorophyll *a*. This similarity suggests that specific LhcE proteins, LhcE5–8, LhcE10–11, and LhcE13, contribute to the far-red absorption features of the PSI–LHC supercomplex, while LhcE1–4 and LhcE12 are primarily responsible for far-red absorption in the free 180 kDa LhcE antenna complex.

Similar red-shifted antenna features have been described in flowering plants and in green microalgae such as *Chlamydomonas reinhardtii*, where red chlorophylls are predominantly located in LHCI subunits (Croce et al., 1998; Croce et al., 2002; Gibasiewicz et al., 2005). Far-red absorption mechanisms have also been observed in other microalgal lineages, though they involve distinct protein–pigment architectures. For example, in diatoms such as *Phaeodactylum tricornutum*, far-red absorption arises from the oligomerization of a single fucoxanthin–chlorophyll a/c-binding protein, Lhcf15 (Herbstová et al., 2017). Similarly, in the alveolate *Chromera velia*, red-shifted absorption has been reported as well, likely resulting from aggregation of LHC antenna proteins (Bína et al., 2014). Together, these comparisons highlight that although far-red absorption is a widespread adaptation among diverse photosynthetic eukaryotic lineages, *Euglena gracilis* achieves this through a unique set of LHCE proteins and their complex-specific distribution, representing a distinct evolutionary strategy within secondary green plastids.

### 3.6 Dynamic Interaction of the LhcE Antenna Complex with PSII

In photosynthetic organisms, the ability to dynamically adjust antenna size and composition is key to optimizing light use efficiency. We therefore investigated how the 180 kDa LhcE antenna complex responds to changing light conditions in *E. gracilis*, focusing on both long-term high-light acclimation and short-term far-red light exposure.

Comparison of pigment–protein complex distributions under varying light conditions (from high-light to very low light) revealed a marked reduction in the 180 kDa LhcE antenna complex under high-light (HL) regimes (Figure 6A). This decline was accompanied by a measurable decrease in far-red absorption capacity in whole cells (Figure 6B). Additionally, HL-acclimated cells exhibited (i) reduced fluorescence emission at 705 nm at room temperature (Figure 6C), and (ii) a significant decrease in the chlorophyll *a/b* ratio (from 6.2 ± 0.2 in low light (LL) to 5.1 ± 0.3 in HL; *p* < 0.005), a rare trend among photosynthetic organisms (Nagao et al., 2021). Together, these findings suggest that modulation of LhcE antenna complex abundance represents a primary long-term acclimation strategy to light intensity in *E. gracilis*.

**Figure 6.**
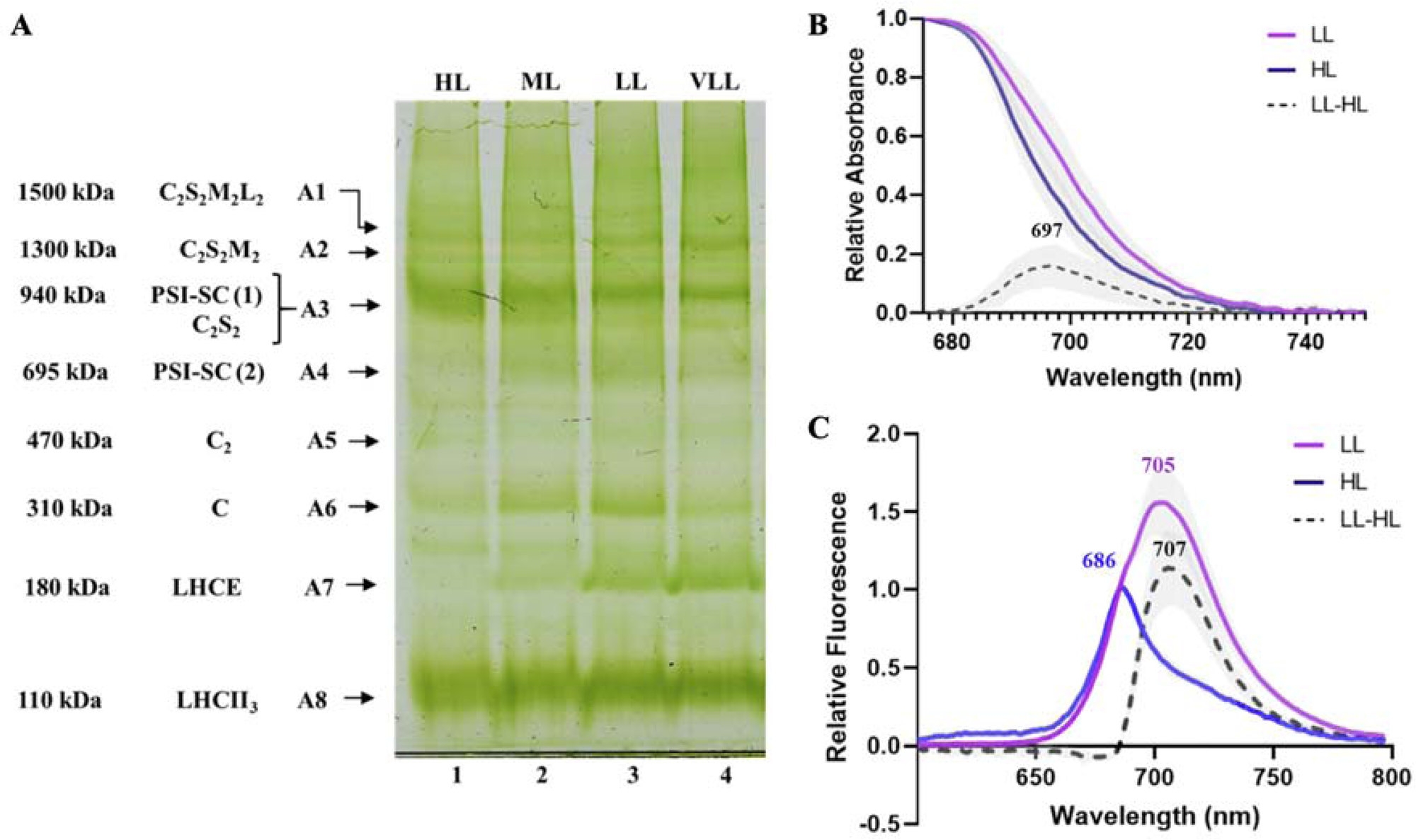
Dynamics of LCHE antenna complex under changing light conditions. **A.** *hr*CN-PAGE of photosynthetic pigment-proteins complexes in *E. gracilis* cells grown in different light intensities (high light (HL), medium light (ML), low light (LL), very low light (VLL). **B.** Absorption spectra of whole cells grown in LL and HL. **C.** Room temperature fluorescence spectra of whole cells grown in LL and HL. Values are represented as mean ± s.d. (n=3, based on independent biological replicates).

In short-term experiments, *E. gracilis* cells acclimated to low light (LL) were exposed to far-red light (720 nm), resulting in a transient increase in PSII maximum fluorescence at room temperature (146 ± 9% relative to baseline), which returned to initial levels after dark acclimation (Figure 7A). The fluorescence emission difference spectrum between far-red and dark-acclimated cells peaked at 707 nm (Figure 7B), closely resembling the emission profile of the isolated 180 kDa LhcE antenna complex (Figure 5B). A similar red-shifted spectral component was also observed in the emission spectra of LL-versus HL-acclimated cells (Figure 6C). Taken together, these data indicate that in LL-acclimated cells, short-term far-red exposure triggers the reversible association of the LhcE antenna complex with PSII. This interpretation is further supported by a significant increase in PSII absorption at 720 nm (far-red) relative to 660 nm (red) in far-red–acclimated cells compared to dark-acclimated controls (Figure 7C). In contrast, cells grown under high light, where the 180 kDa LhcE complex is nearly undetectable (Figure 6A) and far-red fluorescence emission is strongly suppressed (Figure 6C), showed no significant change in PSII antenna characteristics between far-red and dark acclimation states (Figure 7C and 7D).

**Figure 7.**
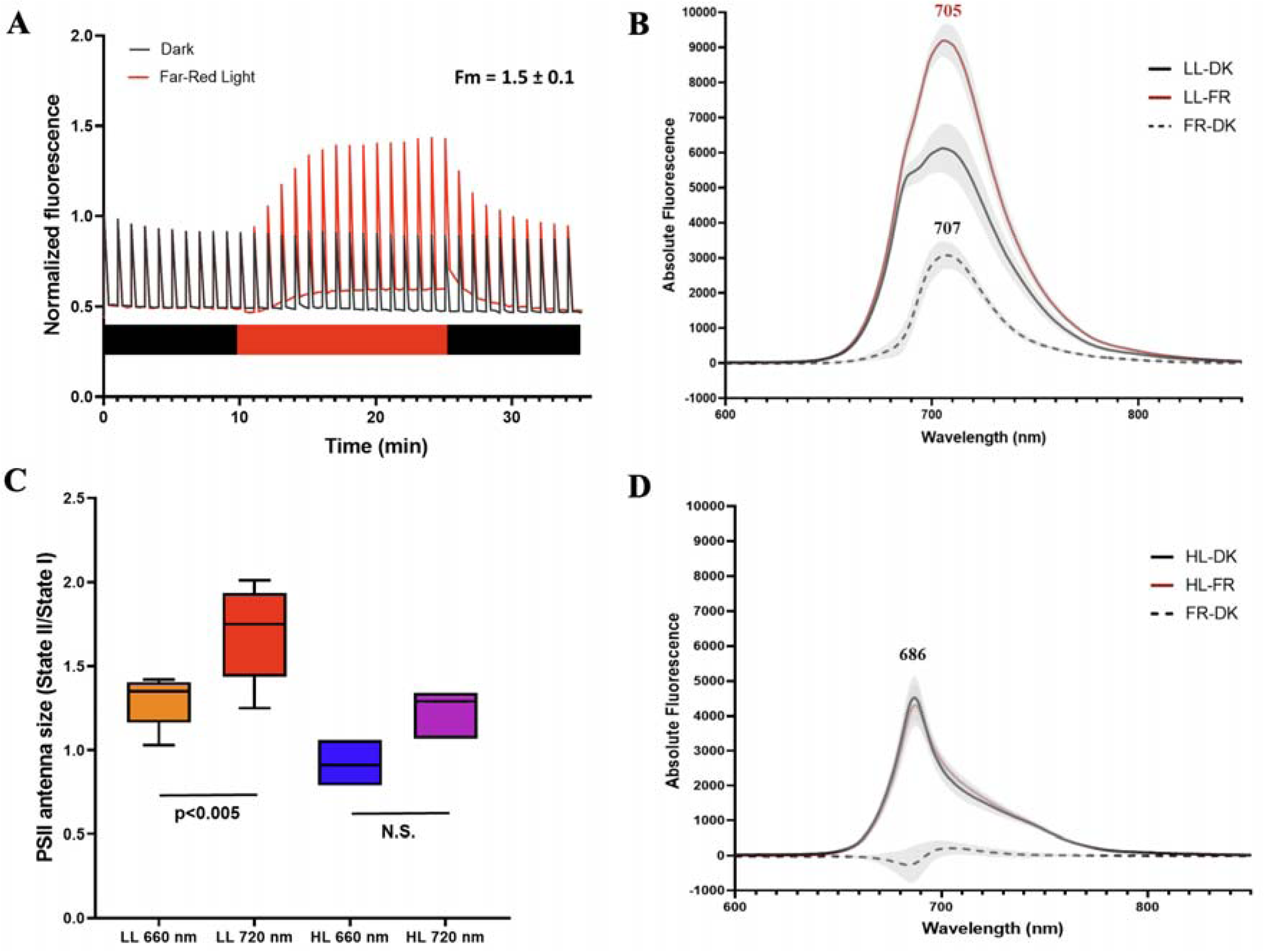
LhcE antenna complex dynamic interaction with PSII. **A.** *In vivo* monitoring of maximum chlorophyll a fluorescence was performed on LL-grown cells in darkness (blue line), or during 15-min exposure to FR (720 nm) illumination followed by 10-min dark recovery period (red line). The red/black bar indicates the period of FR illumination and dark, respectively. **C.** PSII antenna size was estimated upon illumination at 660 nm or 720 nm after exposure of LL and HL cells to darkness (state II) or to FR (state I). **B,D.** Room Temperature fluorescence spectra of cells after exposure to darkness (DK) or far-red illumination (FR) for 15 minutes. LL and HL indicate the light intensity of growing conditions, 25 and 450 PPFD, respectively (n=3, based on independent biological replicates).

## Discussion

Chlorophyll a is common to almost all oxygenic photosynthetic organisms. In rhodophytes, glaucophytes, and most cyanobacteria, it is the only chlorophyll species functioning in the photosystems (Gould et al., 2008). In contrast, land plants and green algae also rely on chlorophyll *b* to stabilize major LHC complexes, although chlorophyll *a* remains essential for photochemistry (Tanaka and Tanaka, 2011). The newly proposed LhcE protein family in *E. gracilis* contains only chlorophyll *a* and diadinoxanthin (Table 1), which contributes to the characteristically low chlorophyll *b* content in this species (Cunningham and Schiff, 1986). Phylogenetically, this large family is distinct from Viridiplantae LHC proteins (Figure 1B), suggesting it expanded independently from a small number of ancestral genes following chloroplast acquisition from a *Pyramimonas*-like green alga (Turmel et al., 2009). Members of the LhcE family are found in both the PSI–LHC supercomplex and the free 180 kDa antenna complex, although LhcE9 also associates with PSII. The 180 kDa LhcE complex is likely organized as a pentamer containing LhcE1–4 and LhcE12 (Figure 5), a structure not previously observed in green plastids (Iwai et al., 2024). Interestingly, a similar tetramer/pentamer configuration of fucoxanthin chlorophyll *a/c*-binding proteins was recently described in the diatom *Chaetoceros gracilis* (Zhou et al., 2024), suggesting potential functional convergence among photosynthetic eukaryotes. Beyond its unique pigment composition, the *E. gracilis* LhcE antenna complex exhibits several remarkable features. Its absorption spectrum shows a red-shifted secondary peak (∼698 nm), approximately 20 nm beyond that of LhcbM trimers (Figure 5A).

This LhcE antenna complex also appears highly dynamic: its abundance decreases under increasing light intensities (Figure 6), in parallel with an increase in the chlorophyll *a/b* ratio, indicating a role in acclimation to light conditions. This flexibility may reflect adaptation to variable light environments (Bag, 2021) or an ancestral origin. The latter hypothesis is supported by the deep phylogenetic branching of LhcE proteins and the absence of canonical LHCA genes in our phylogenetic analysis (Figure 1B; Supplementary Fig. S48), consistent with a limited LHC gene diversity at the origin of euglenid photosynthetic lineages.

It has been proposed that early photosynthetic eukaryotes used LHC proteins to supply PSI, while PSII light capture was supported by phycobilisomes. Over evolutionary time, green algae and land plants developed more specialized photosystem-specific LHCs, with LHCII associating almost exclusively with PSII (Pan et al., 2020). This PSII-specific association is consistent with the C□S□M□L□-type PSII SC we observed in *E. gracilis* (Figure 3C).

Structural analyses reveal many Lhc proteins associated with *E. gracilis* PSI, including LhcbM2, LhcbM8, and at least ten LhcE proteins (Figure 4A), far exceeding the PSI antenna sizes in green lineage species (Gorski et al., 2022; Qin et al., 2019). Remarkably, LhcbM2 and LhcbM8 form basal branches in the Lhcb phylogenetic tree (Figure 1), suggesting they evolved independently of the PSII-associated LhcbM proteins found in Viridiplantae. A comparable PSI antenna expansion is seen in other secondary photosynthetic lineages, including the PSI–FCPI supercomplex in *Chaetoceros neogracilis* (Xu et al., 2020) and the PSI–LHC complex in the symbiotic dinoflagellate *Symbiodinium* (Zhao et al., 2024). Similarly, a PSI antenna expansion is observed as a reversible photoacclimation to low-light which leads to the formation of a PSI-LHCI-Lchp supercomplex in prasinophyte species *Ostreococcus tauri* (Ishii et al., 2023). In *Euglena*, this expanded PSI antenna is associated with a minimal PSI core containing just eight subunits, PsaA-F/J/M (Supplementary Table S3). The absence of most peripheral PSI subunits appears to reflect convergent evolution, common to several secondary plastid-containing lineages (Basso et al., 2014; Neilson et al., 2017; Sobotka et al., 2017).

In Viridiplantae, LHC function is modulated through adaptive mechanisms such as NPQ and state transitions, which dynamically redistribute excitation energy captured by LHCII between PSII and PSI (Virtanen and Tyystjärvi, 2023). State transitions are driven by the redox state of the plastoquinone pool and involve phosphorylation of loosely or free LHCII□ complex, which migrates from PSII region of the thylakoid membrane and rebinds to PSI (Longoni et al., 2019; Huang et al., 2021; Pan et al., 2021). In contrast, our data indicate that a canonical state transition mechanism involving LhcbM trimers is absent in *E. gracilis*. Instead, we observed that in LL–acclimated cells, short-term exposure to far-red light induces dynamic association of the pentameric LhcE antenna complex with PSII. This light-triggered interaction is supported by red-shifted fluorescence emission and enhanced far-red absorption, suggesting a previously unrecognized, state transition–like mechanism. This is reminiscent of the hypothesis proposed as early as 1961 by Brown and French that *E. gracilis* uses a common antenna system. Our findings support this idea, particularly with respect to the species-specific far-red–emitting LhcE complex (Figures 1C and 4E). Although CP29 is known to mediate state transitions in *C. reinhardtii* (Tokutsu et al., 2009), its role in *Euglena* appears limited, given that it is surrounded by LhcbM trimers which do not dissociate (Figure 3). Instead, we propose that LhcE9, which is consistently found in all PSII SCs (Figure 3A), may provide a structural interface for docking the 180 kDa LhcE antenna complex. The absence of CP26 (Figure 3C), a conserved minor PSII antenna in green algae and land plants, may also be linked to the acquisition of this novel LhcE-based regulatory antenna system.

The *Euglena* PSI core is surrounded by a belt of LhcE and LhcbM proteins (Figure 4C), a configuration that likely constrains canonical state transition mechanisms. Moreover, in *E. gracilis,* the absence of PsaH, PsaL, and PsaO, key subunits required for LHCII□ binding in Viridiplantae (Yang et al., 2015), further supports this divergence. Instead, the reversible association of the red-shifted 180 kDa LhcE complex with PSII (Figure 7) appears to function as an alternative mechanism for optimizing light harvesting under low-light or far-red–enriched conditions.

## Concluding Remarks

Light adaptation mechanisms are essential for the survival of photosynthetic organisms under fluctuating illumination (Finazzi et al., 1999). As described above, most classical acclimation strategies observed in Viridiplantae appear to be either lost or were never acquired by *Euglena gracilis* during green plastid endosymbiosis. Instead, *E. gracilis* has evolved a lineage-specific, mobile light-harvesting antenna, designated here as LhcE, which plays a central role in light acclimation. This LhcE antenna complex, together with PSI-associated LhcE proteins, is responsible for the far-red absorption capacity of *E. gracilis* cells. It accumulates under low-light and far-red conditions, enhancing photon capture when light availability is limited. More than sixty years after the first observations of light-dependent spectral shifts in *E. gracilis* (Brown and French, 1961), we have identified the specific antenna complex responsible for this phenomenon. Remarkably, this unique LhcE complex, not the classical LHCII□ antenna, is dynamically involved in a state transition-like mechanism in *E. gracilis*, highlighting a distinct strategy for balancing excitation energy in this lineage. Overall, our findings reveal a fundamentally different organization of the photosynthetic apparatus in *E. gracilis* that promotes adaptive light harvesting through a previously unrecognized mechanism. A model summarizing these features and their functional implications is presented in Figure 8.

**Figure 8.**
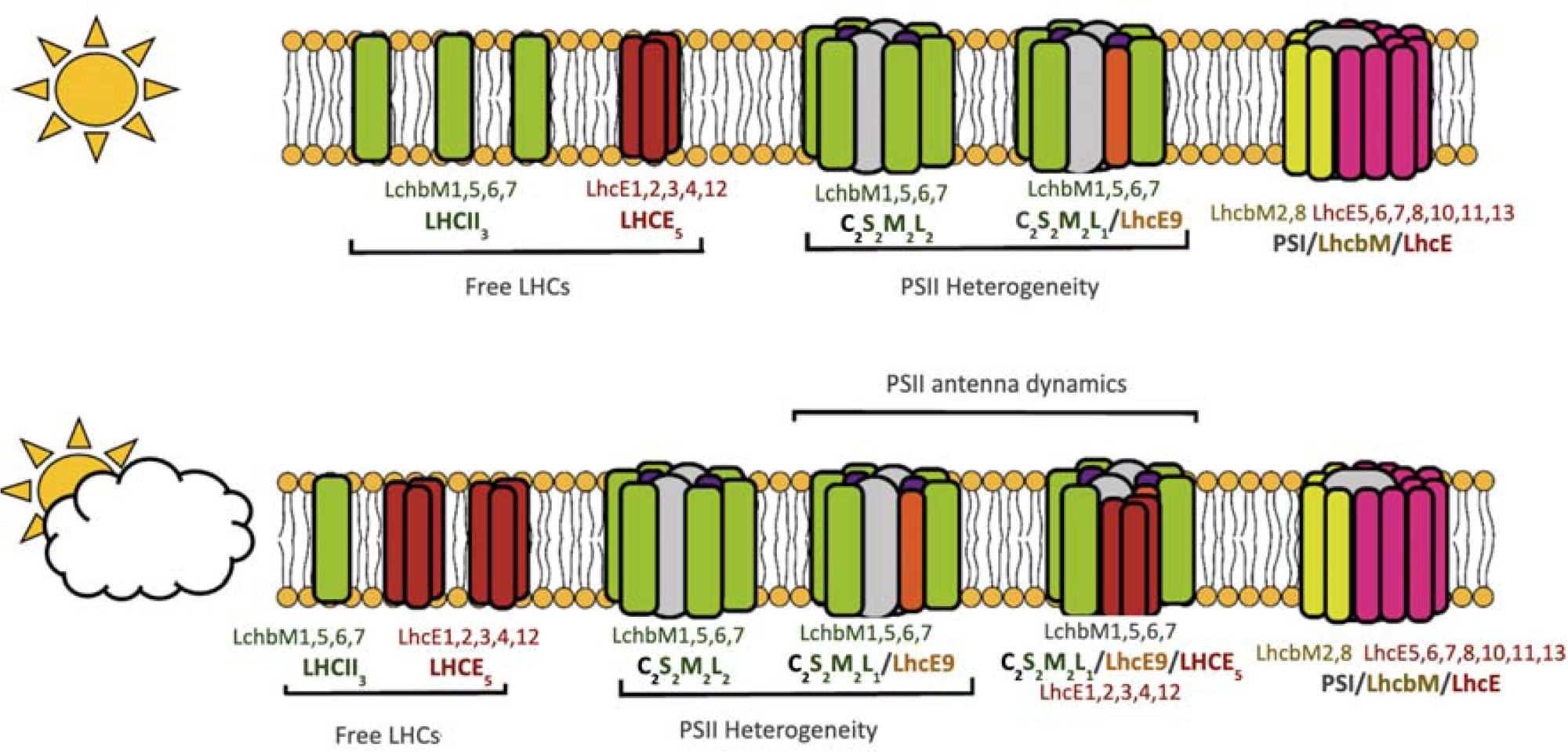
Proposed mechanism for light adaptation in *Euglena gracilis*. Diversity of photosynthetic pigment-proteins complexes distribution among thylakoid membranes depending on light fluctuations: *upper panel*: medium light; *lower panel*: low light. Free antennas (LHCII_3_ and LHCE_5_), PSII heterogeneity (C_2_S_2_M_2_L_2_/C_2_M_2_L_1_LHCE9), PSII/LHCE_5_ dynamic regulation and PSI-LHCE-LHCBM supercomplex are indicated.

## Supplementary data

**The following supplementary data are available at Zenodo.**

https://doi.org/10.5281/zenodo.15353545

Table S1. Corresponding nomenclature for *Euglena gracilis* LHC sequences.

Table S2. Quantitative Proteomic Analysis of Representative *Euglena gracilis* samples

Table S3. Composition of the photosystem I and II cores complex in *Euglena gracilis*.

Fig. S1-47. Phylogenetic trees of *Euglena gracilis* PSI/PSII subunits.

Fig. S48. Phylogenetic tree of *Euglena gracilis* Light-Harvesting Complex (Lhc) Proteins.

Fig. S49. Structural comparison of Light Harvesting Complexes proteins.

Fig. S50. Estimated molecular masses for the *Euglena gracilis* photosynthetic complexes.

Fig. S51. Distribution of LhcbM in PSII-SC and free LHCII across different PSII/LHCII particles.

Fig. S52. Projection maps and structural models of PSII supercomplexes from *Euglena gracilis* revealed by single particle electron microscopy.

Fig. S53. Structural models of PSI-LHC supercomplexes from *Euglena gracilis* revealed by single-particle electron microscopy.

Dataset S1. Reference amino acid sequences for LHC, PSI, and PSII subunits compiled from the literature for model organisms.

Dataset S2. Comprehensive set of 158 predicted Lhc protein sequences in *E. gracilis*.

Dataset S3. PSI and PSII subunit amino acid sequences in *E. gracilis*.

## Acknowledgements

We thank Paulina Karpinska for technical assistance and Roberta Croce for her valuable scientific advice.

## Author contributions

H.M.A. and P.C.: conceptualization; H.M.A., W.N., H.D., D.B., R.K. and P.C.: methodology; H.M.A., R.A., W.N., D.B., R.K. and P.C.: formal analysis; H.M.A., R.A., F.V.L., Z.A.G., H.F., T.F., A.G., W.N., C.C. and H.D..: investigation; H.M.A., P.M., D.B., R.K. and P.C.: resources; H.M.A., R.A., T.F., P.M., D.B., R.K. and P.C.: data curation; H.M.A. and P.C.: writing - original draft; H.M.A., R.A., F.V.L., T.F., W.N., D.B., R.K. and P.C.: writing - review & editing; H.M.A., R.A., D.B., R.K. and P.C.: visualization; H.M.A., D.B., R.K. and P.C.: supervision; H.M.A., D.B., R.K. and P.C.: funding acquisition

## Data availability

All primary data supporting the results of this study are available from the corresponding author upon reasonable request. This includes raw data used to generate graphs, spectra, and histograms.

## Abbreviations

α-DDM: n-dodecyl-α-D-maltoside
β-DDM: n-dodecyl-β-D-maltoside
CC: Core complex
DCMU: 3-(3,4-dichlorophenyl)-1,1-dimethylurea
HL: High light
hrCN-PAGE: high resolution clear native PAGE
kDa: kiloDalton
ML: Medium light
LC-ESI-Q-TOF: Liquid chromatography–electrospray ionization–quadrupole time-of-flight
LHC: Light Harvesting complex
LL: Low Light
µE: µmol photons
NPQ: Non photochemical quenching
PPFD: Photosynthetic photon flux density
RC: Reaction center
VLL: Very low light

